# Carbon content, carbon fixation yield and dissolved organic carbon release from diverse marine nitrifiers

**DOI:** 10.1101/2022.01.04.474793

**Authors:** Barbara Bayer, Kelsey McBeain, Craig A. Carlson, Alyson E. Santoro

## Abstract

Nitrifying microorganisms, including ammonia-oxidizing archaea, ammonia-oxidizing bacteria and nitrite-oxidizing bacteria, are the most abundant chemoautotrophs in the ocean and play an important role in the global carbon cycle by fixing dissolved inorganic carbon (DIC) into biomass. The release of organic compounds by these microbes is less well known but may represent an as-yet unaccounted source of dissolved organic carbon (DOC) available to heterotrophic marine food webs. Here, we provide measurements of cellular carbon and nitrogen quotas, DIC fixation yields and DOC release of ten phylogenetically diverse marine nitrifiers grown in multiple culture conditions. All investigated strains released DOC during growth, making up on average 5-15% of the fixed DIC. Neither substrate concentration nor temperature affected the proportion of fixed DIC released as DOC, but release rates varied between closely related species. Our results also indicate previous studies may have underestimated DIC fixation yields of marine nitrite oxidizers due to partial decoupling of nitrite oxidation from CO_2_ fixation, and due to lower observed yields in artificial compared to natural seawater medium. The results of this study provide values for biogeochemical models of the global carbon cycle, and help to further constrain the implications of nitrification-fueled chemoautotrophy for marine food-web functioning and the biological sequestration of carbon in the ocean.

## Introduction

Marine microorganisms play a critical role in the global carbon cycle through their transformations of organic and inorganic carbon constituents. A fraction of the carbon dioxide (CO_2_) that is captured by phytoplankton in the surface ocean sinks to depth as dead organic material, supporting a mesopelagic food web of both microbes and higher trophic levels (Hannides et al. 2013; Giering et al. 2014; Choy et al. 2015). Organic matter decomposition in the mesopelagic also releases ammonium, a reduced form of nitrogen that can be used as an energy source by chemoautotrophic nitrifying archaea and bacteria to fuel dissolved inorganic carbon (DIC) fixation into biomass (Ward 2011). Chemoautotrophic production provides a new, labile, non-sinking source of particulate organic matter to the deep ocean which is otherwise dominated by refractory organic carbon (Reinthaler et al. 2010; Middelburg 2011), supporting a significant fraction of the heterotrophic microbial community in the mesopelagic (Hansman et al. 2009).

The main nitrifiers in the ocean are ammonia-oxidizing archaea (AOA), which oxidize ammonia (NH_3_) to nitrite (NO_2_^−^), and nitrite-oxidizing bacteria (NOB), which further oxidize NO_2_^−^ to nitrate (NO_3_^−^) (Ward 2011). These two steps are assumed to be tightly coupled, as NO_2_^−^ typically does not accumulate in oxic open ocean waters (with the exception of the primary nitrite maximum at the base of the euphotic zone (Lomas and Lipschultz 2006; Santoro et al. 2013)). Despite this tight coupling, AOA are approximately six times more abundant than NOB at a given location and sampling depth (Santoro et al. 2019), possibly owing to their smaller cell size compared to NOB (Watson and Waterbury 1971; Könneke et al. 2005; Santoro and Casciotti 2011; Bayer et al. 2016; Mueller et al. 2021), or as a result of the higher theoretical energy yield from ammonia compared to nitrite oxidation (Bock and Wagner 2013). Ammonia-oxidizing bacteria (AOB) are thought to play a minor role in global ocean nitrification due to their overall low abundances (Santoro et al. 2010; Buchwald et al. 2015; Tolar et al. 2016).

Despite the known difference in theoretical energy yield, there are many uncertainties regarding the organic carbon yield from ammonia versus nitrite oxidation (hereinafter referred to as DIC fixation yield) and the contribution of both functional groups to chemoautotrophic DIC fixation in the ocean. AOA cultures have recently been shown to release dissolved organic carbon (DOC) during growth (Bayer et al. 2019a), pointing to a potential loss of cellular fixed carbon that is not captured by conventional methods measuring DIC incorporation into biomass. The release of DOC by nitrifiers might represent an as-yet unaccounted source of organic material in the deep ocean potentially fueling the microbial loop, with important implications for the marine carbon cycle. However, it remains unclear if DOC release is a phenomenon only observed under specific culture conditions restricted to some AOA, or a common feature shared by diverse autotrophic nitrifiers under natural conditions.

Here, we report combined measurements of DIC fixation and DOC release of ten phylogenetically diverse marine nitrifiers comprising two AOA genera (*Nitrosopumilus* and *Ca*. Nitrosopelagicus), one AOB genus (*Nitrosomonas*) and three NOB genera (*Nitrospina, Nitrospira*, and *Nitrococcus*), and further explore the effect of substrate concentration, temperature, and different culture media on these measurements. The results of this study will inform ecological theoretical models to further constrain DIC fixation yields associated with nitrification in order to better understand the dynamics involved in the sequestration of carbon in the ocean.

## Methods

### Nitrifier culture sources

The AOA cultures used in this study were three axenic *Nitrosopumilus* strains and one *Nitrosopelagicus* enrichment culture. *Ca*. Nitrosopelagicus brevis U25 originates from a North Pacific Ocean water sample (Santoro and Casciotti 2011; Carini et al. 2018). The level of enrichment during the time of this study was ∼90%. *Nitrosopumilus* sp. CCS1 is a novel AOA strain isolated from a seawater sample collected from the California Current system in the North Pacific Ocean (Santoro et al, unpublished). *Nitrosopumilus adriaticus* NF5 (=JCM 32270^T^ =NCIMB 15114^T^) and *Nitrosopumilus piranensis* D3C (=JCM 32271^T^ =DSM 106147^T^ =NCIMB 15115^T^) were isolated from the Northern Adriatic Sea and have been described in detail (Bayer et al. 2016, Bayer 2019).

The four axenic NOB strains, *Nitrospina gracilis* Nb-211, *Nitrospina* sp. Nb-3, *Nitrococcus mobilis* Nb-231 and *Nitrospira marina* Nb-295, were obtained from the culture collection of John B. Waterbury and Frederica Valois at the Woods Hole Oceanographic Institution (WHOI). *N. gracilis* Nb-211 was isolated from surface waters of the South Atlantic Ocean (Watson and Waterbury 1971), *N. mobilis* Nb-231 was isolated from a surface water sample obtained from the South Pacific Ocean (Watson and Waterbury 1971) and *N. marina* Nb-295 was isolated from a water sample collected at a depth of 206 m from the Gulf of Maine in the Atlantic Ocean (Watson et al. 1986). *Nitrospina* sp. Nb-3 was isolated from the Pacific Ocean off the coast of Peru and has not yet been officially described (Watson and Waterbury, unpublished), however, it shares a high 16S rRNA gene sequence similarity with strain 3/211 (Lücker et al. 2013).

AOB strains used in this study, *Nitrosomonas marina* C-25 and *Nitrosomonas* sp. C-15 (also known as strain Nm51, (Koops et al. 1991)), were both obtained from the culture collection of John B. Waterbury and Frederica Valois at WHOI and were revived from 60-year old cryostocks. Strain C-15 was isolated from surface water (1 m depth) of the South Pacific Ocean off the Peruvian continental shelf (Watson and Mandel 1971) and strain C-25 was isolated from surface waters of the South Atlantic Ocean (200 miles off the Amazon River mouth) (Watson and Mandel 1971).

### Culture conditions

*Nitrosopumilus adriaticus* NF5, *Nitrosopumilus piranensis* D3C, *Nitrosomonas marina* C-25 and *Nitrosomonas* sp. C-15 were grown in HEPES-buffered artificial seawater medium containing 1 mM NH_4_Cl, and *Ca*. Nitrosopelagicus brevis U25 was grown in natural seawater medium containing 50 µM NH_4_Cl. *Nitrospina gracilis* Nb-211, *Nitrospira marina* Nb-295 and *Nitrococcus mobilis* Nb-231 were grown in artificial seawater medium supplemented with 1 mM NaNO_2_. *Nitrosopumilus* sp. CCS1 and *Nitrospina sp*. Nb-3 were grown under multiple culture conditions as indicated in the Results and Discussion. All strains were routinely grown in 60 mL polycarbonate bottles (Nalgene) containing 50 mL culture medium, and bottles were incubated at either 15°C or 25 °C (with the exception of *Ca*. Nitrosopelagicus brevis which was always incubated at 22°C) in the dark without agitation.

The artificial seawater medium contained 18.54 g L^-1^ NaCl, 4.7 g L^-1^ MgSO_4_ × 7H_2_O, 3.55 g L^-1^ MgCl_2_ × 6H_2_O, 1.03 g L^-1^ CaCl_2_ × 2H_2_O, 0.51 g L^-1^ KCl, 0.14 g L^-1^ NaHCO_3_. The natural seawater medium consisted of aged seawater collected from the Santa Barbara Channel (approx. 10 m depth at, 0.2 μm pore size filtered). Artificial and natural seawater were supplemented with 2.6 mg L^-1^ K_2_HPO_4_, 250 µg L^-1^ FeNaEDTA, 30 µg L^-1^ H_3_BO_3_, 20µg L^-1^ MnCl_2_ × 4H_2_O, 20 µg L^-1^ CoCl_2_ × 6H_2_O, 24 µg L^-1^ NiCl_2_ × 6H_2_O, 20 µg L^-1^ CuCl_2_ × 2H_2_O, 144 µg L^-1^ ZnSO_4_ × 7H_2_O, 24 µg L^-1^ Na_2_MoO_4_ × 2H_2_O. The pH was adjusted to 7.8-8.0 with NaOH or HCl. Due to the pH decrease associated with ammonia oxidation, culture medium with high initial NH_4_^+^ concentrations (>250 µM) was buffered by addition of 10 mM HEPES (pH 7.8). AOA cultures were supplemented with 50 U L^-1^ catalase (Sigma-Aldrich, Cat. Nr. C9322) to reduce oxidative stress and NOB cultures were supplemented with 50 ng L^-1^ cyanocobalamin. To test the effect of reduced and organic nitrogen compounds on *Nitrospina sp*. Nb-3, NH_4_Cl (50 µM) or tryptone (150 mg L^−1^) were added to the culture medium.

NO_2_^−^ concentrations were measured using the Griess-Ilosvay colorimetric method (Strickland and Parsons 1972) and enumeration of cells was performed on an Easy-Cyte flow cytometer (Guava Technologies) following SYBR Green staining as previously described (Bayer et al. 2021).

### Cellular carbon and nitrogen content measurements

To determine C : N ratios, ca. 100-500 mL of culture was filtered onto combusted (450°C, 4h) glass fiber filters (Advantec, GF-75, 25mm). Filters were acidified with HCl (10% v/v), dried (60°C, 24 h), and packed into tin capsules prior to being analyzed on a CHN elemental analyzer (Exeter Analytical, CEC 440HA). The instrument was calibrated with acetanilide following manufacturer protocols.

Cellular carbon (C) content was calculated using both, CHN elemental analyzer (only for large cells) and ^14^C-DIC incorporation measurements (see below), divided by the number of newly produced cells. Additionally, C content of the AOA strain *Nitrosopumilus* sp. CCS1 was calculated from a dilution series of concentrated cells as described in (White et al. 2019). Cells were concentrated using tangential flow filtration (Pellicon) and a dilution series of 1.1 to 5.6 ×10^11^ cells L^-1^ was constructed by resuspending cell concentrates in culture medium (Fig. S1). The total organic C content for each vial of the dilution series was directly measured by high temperature combustion using a modified Shimadzu TOC-V as described in (Carlson et al. 2010). C content per cell was calculated via linear regression of cell counts and elemental content over the dilution series, where the slope of a Model II least squares regression is considered the elemental content per cell (Fig. S1).

### Combined DIC fixation and DOC release measurements

DIC fixation was measured via the incorporation of [^14^C]-bicarbonate as previously described (Herndl et al. 2005) with modifications. [^14^C]-bicarbonate (specific activity 56 mCi mmol^-1^/2.072 × 10^9^ Bq mmol^-1^, Perkin Elmer) was added to 5 mL of culture (between 10-60 µCi were added depending on the activity of the culture). Different incubation times were tested (see Results section) and all consecutive experiments were performed over the entire length of the growth curve. For every culture condition, at least three replicate live samples and one formaldehyde-fixed blank (3% v/v) were incubated in temperature-controlled incubators in the dark. Parallel incubations without [^14^C]-tracer additions were used to determine cell abundance and nitrite concentration (see above).

Incubations were terminated by adding formaldehyde (3% v/v) to 5 mL of sample. After 30-60 min, every sample was individually filtered onto 25 mm, 0.2 µm pore size polycarbonate filters (Millipore) and rinsed with 0.5 mL of artificial seawater using a glass filtration set (Millipore). The individual filtrates (5.5 mL per sample) were collected and transferred to scintillation vials to determine the fraction of [^14^C]-dissolved organic carbon ([^14^C]-DOC). Excess [^14^C]-bicarbonate from the filters was removed by exposing them to fumes of concentrated HCl (37 %) for 24 h. The filters were transferred to scintillation vials and 10 mL of scintillation cocktail (Ultima Gold, Perkin Elmer) was added. The filtrates were acidified to pH ∼2 with HCl (25 %) as previously described (Marañón et al. 2004), and filtrates were kept for 24 h in open scintillation vials placed on an orbital shaker before 10 mL scintillation cocktail was added to each vial. Samples were shaken for ca. 30 sec and incubated in the dark for at least 24 hours prior to counting the disintegrations per minute (DPM) in a scintillation counter (Beckman Coulter LS6500) for 15 min.

Total radioactivity measurements were performed to verify added [^14^C]-bicarbonate concentrations by pipetting 100µl of sample into scintillation vials containing 400µl beta-phenylethylamine (to prevent outgassing of ^14^CO_2_). Scintillation cocktail was added, vials were shaken for ca. 30 sec and immediately measured in the scintillation counter.

The resulting mean DPM of the samples were corrected for the DPM of the blank, converted into organic carbon fixed over time and corrected for the DIC concentration in the culture media.

DIC fixation rates were calculated using the following formula:

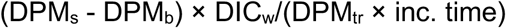

where DPM are the disintegrations per minute measured in the scintillation counter, for the sample (s) and the blank (b). DIC_w_ denotes the dissolved inorganic carbon concentration in culture medium and DPM tracer (tr) is the DPM for the [^14^C]-bicarbonate added to the incubations.

### DIC concentration measurements

Total alkalinity (TA) of unfixed natural and artificial seawater medium was measured via an open-cell endpoint titration using a Mettler-Toledo T5 autotitrator, and pH was measured spectrophotometrically using a Shimadzu UV-1280 UV-VIS spectrophotometer as described previously (Dickson et al. 2007; Hoshijima and Hofmann 2019). Dissolved inorganic carbon (DIC) concentrations were calculated from TA and pH using the CO2SYS software (Pelletier 2007). To calculate DIC concentrations of HEPES-buffered media, TA values were taken from unbuffered artificial seawater medium and the pH was re-measured after adding HEPES.

### *Calculations of Gibbs free energy* (ΔG)

The effective Gibbs free energy (Δ*G*) for ammonia and nitrite oxidation was calculated for the culture conditions in this study using the following formula:

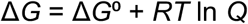

where *R* is the ideal gas constant (8.314 J mol^-1^ K), *Q* is the reaction quotient, and *T* is the temperature in Kelvin. Δ*G*° values were obtained from (Amend and Shock 2001).

*Q* was calculated based on the following measurements and estimates: NO_2_^-^ concentrations were measured directly (see above); [NO_3_^-^] and [NH_4_^+^] were estimated from the decrease or increase in [NO_2_^-^], respectively; NH_3_ concentrations were calculated based on [NH_4_^+^], pH of the culture medium, and the acid association constant (pKa = 9.4); and O_2_ concentrations were estimated to be 235 µM under completely oxic conditions during our incubations. A correction for ionic strength was applied according to (Amend and LaRowe 2019). Calculations can be found in the Supporting Information (Table S1).

### Statistical analyses

Pairwise comparisons were performed with a two-sided Mann-Whitney U Test (pairwise.wilcox.test) using the R software environment (R Core Team 2013). *P* values were adjusted for multiple comparisons using the Benjamini-Hochberg correction (p.adjust.method=“fdr”) (Benjamini and Hochberg 1995).

## Results and Discussion

### Elemental composition of cultured nitrifiers

We determined the cellular carbon (C) content of cultured isolates of ammonia-oxidizing archaea (AOA), ammonia-oxidizing bacteria (AOB) and nitrite-oxidizing bacteria (NOB) belonging to six different genera. The cellular C contents of AOA were ∼11-17 fg C cell^-1^ (Table 1), which is slightly higher than values reported for natural populations in the deep Atlantic Ocean (∼8.39 fg cell^-1^, (Herndl et al. 2005)) and an AOA enrichment culture from the Baltic Sea (9 fg cell^-1^, (Berg et al. 2014)), but much lower than values reported for AOA from hypoxic shelf waters of the Gulf of Mexico (50 ± 16 fg cell^−1^, (Kitzinger et al. 2020)). All investigated marine NOB had higher cellular C quotas compared to AOA (Table 1), with *Nitrospina* exhibiting the lowest (∼28-55 fg C cell^-1^) and *Nitrococcus* the highest (∼272-1207 fg C cell^-1^) values (Table 1). The C content of AOA cells remained fairly constant during different growth phases, while C contents of all investigated NOB strains drastically decreased (∼40-70%) from early exponential growth to stationary phase, which was supported by the observation of smaller cells in stationary compared to exponentially growing cultures (data not shown). Cell sizes of natural populations of Nitrospinae bacteria have been reported to be 4-fold (Kitzinger et al. 2020) and 50-fold (Pachiadaki et al. 2017) larger than AOA cells, potentially reflecting these variations in cell size and C content during different growth phases.

**Table 1.**
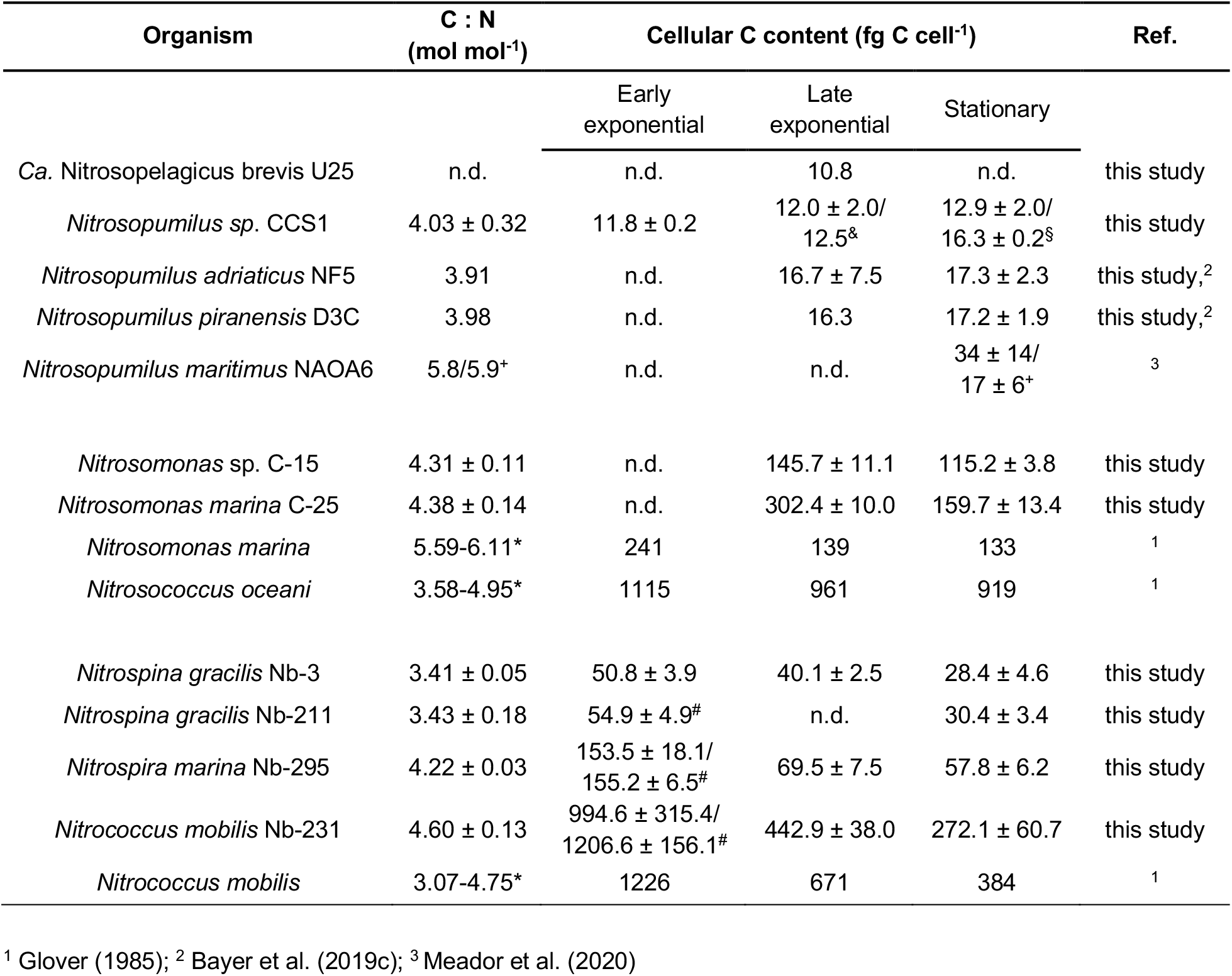
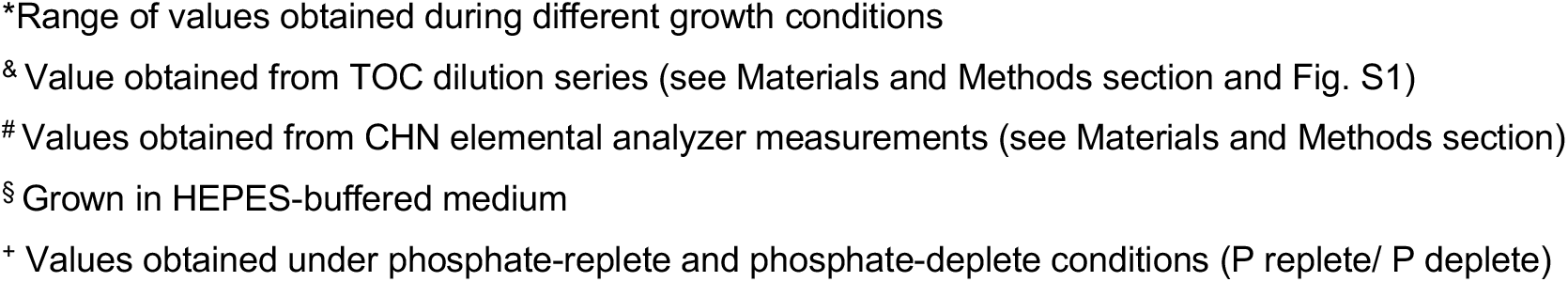
Elemental stoichiometry of phylogenetically diverse cultured marine nitrifiers during different growth phases (early exponential, late exponential, stationary) including previously published values. C : N ratios were obtained during exponential growth phase. Cellular C content values are derived from DIC incorporation measurements if not stated otherwise.

The molar C : N ratios of all investigated nitrifiers were in the range of 3.4-4.6 : 1 (Table 1), with the exception of previously published values of *Nitrosopumilus maritimus* NAOA6 (Meador et al. 2020) and two AOB strains (Glover 1985). The values observed are lower than values of heterotrophic marine bacteria (∼5 : 1) including *Pelagibacter ubique* (∼4.6 : 1) (White et al. 2019), *and references therein*), with *Nitrospina* cells exhibiting the lowest average C : N ratio (∼3.4) of all cultured nitrifiers in our study (Table 1). These low cellular C : N ratios are surprising considering the observation of glycogen storage deposits in cells of *Nitrospina gracilis, Nitrococcus mobilis*, and *Nitrospira marina* (Watson and Waterbury 1971; Watson et al. 1986), as well as polyhydroxbutyrate storage in *Nitrococcus mobilis* (Watson and Waterbury 1971).

### DIC fixation yields of marine nitrifiers

We conducted combined measurements of DIC fixation, DOC release and ammonia/nitrite oxidation rates of ten nitrifier cultures. Here, we use the term ‘DIC fixation yield’ to describe the number of moles of inorganic carbon (CO_2_ or HCO_3_^-^) that are fixed for every mole of N (NH_3_ or NO_2_^-^) oxidized, including the proportion that is released/lost as DOC.

Marine AOA, including three axenic *Nitrosopumilus* strains and one *Ca*. Nitrosopelagicus enrichment culture, exhibited the highest DIC fixation yields (mean±sd= 0.091 ± 0.012, *n*=47) in our study, which were on average ∼2-times higher than those of marine AOB (mean±sd= 0.047 ± 0.010, *n*=23) (Fig. 1). AOA encode the 3-hydroxypropionate/4-hydroxybutyrate (3-HP/4-HB) cycle for DIC fixation (Walker et al. 2010; Santoro et al. 2015; Bayer et al. 2016), which is suggested to be the most energy-efficient aerobic autotrophic DIC fixation pathway, requiring four moles of ATP to fix two moles of C (Könneke et al. 2014). In contrast, AOB use the Calvin-Benson-Bassham (CBB) cycle (Utåker et al. 2002; Stein et al. 2007), which has a higher ATP requirement and an estimated 20% loss of fixed DIC due to the oxygenase side-reaction of ribulose-1,5-bisphosphate carboxylase/oxygenase (Berg 2011). DIC fixation yields of two *Nitrosopumilus* strains were recently reported to be up to ten times higher (0.18-1.2, (Meador et al. 2020)) compared to values in our study and previously published values of *Nitrosopumilus adriaticus* NF5 (0.1, (Bayer et al. 2019c)) and a *Nitrosarchaeum* enrichment culture (0.1, (Berg et al. 2014)). However, such high values would require unrealistically high ATP yields (up to 2.4 moles ATP per mole NH_3_ oxidized) compared to reported estimates of 0.15-0.28 ATP/NH_3_ (mol/ mol) (Li et al. 2018).

**Fig.1.**
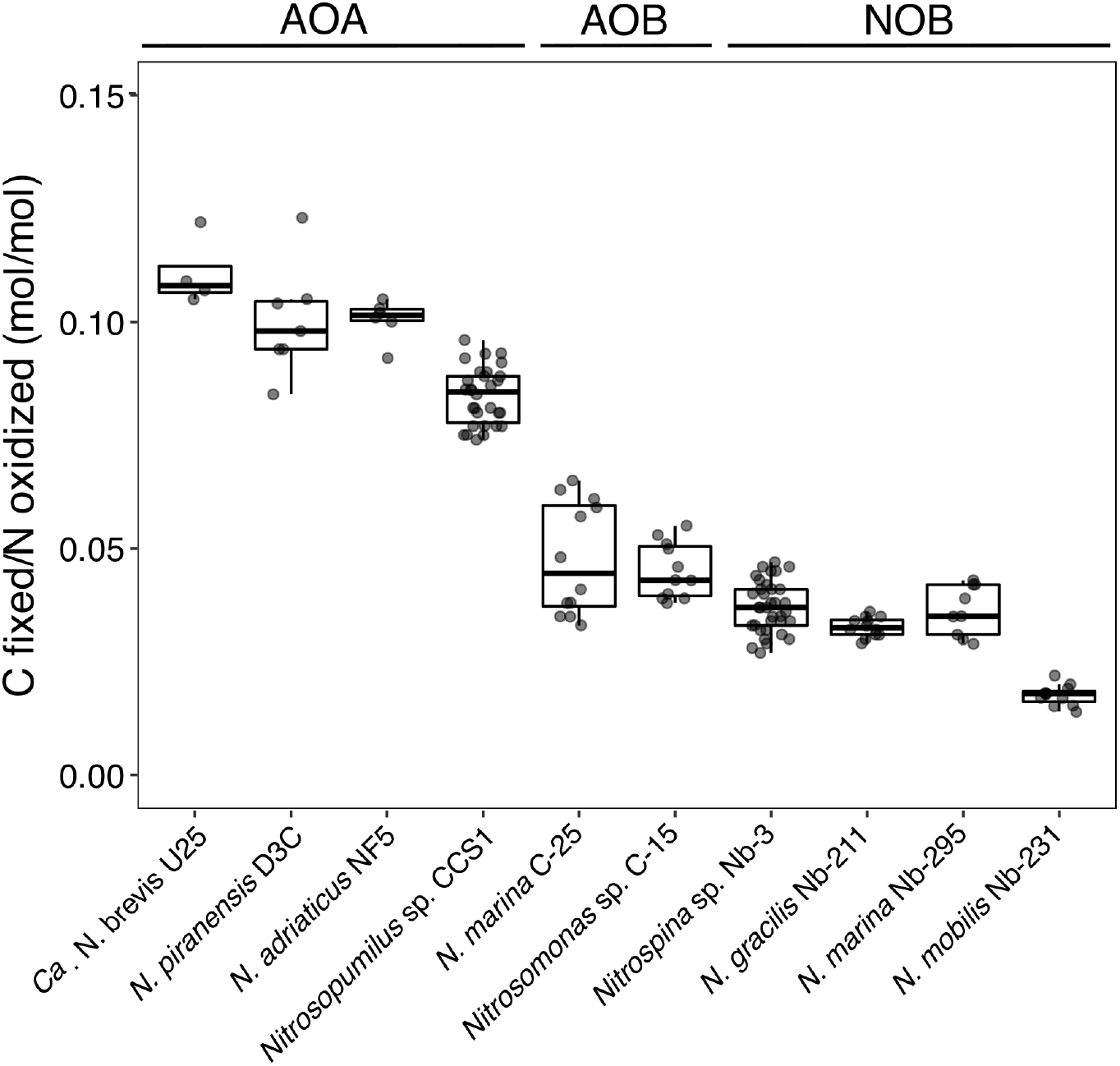
Comparison of DIC fixation yields of ten different phylogenetically diverse marine nitrifiers. Plotted values include both, the fraction of C incorporated into biomass and the fraction of C released as DOC. For NOB, only measurements conducted over the entire length of the growth curve (until stationary phase) are shown (see Fig. 2). Values obtained from cultures grown under different conditions (see Fig. 3 and Fig. S3) are included in this plot.

DIC fixation yields of marine NOB (*Nitrospina/Nitrospira*: mean±sd=0.036 ± 0.005, *n*=47; *Nitrococcus*: mean±sd=0.018 ± 0.002, *n*=11) were lower compared to those of ammonia oxidizers (Fig. 1). *Nitrococcus mobilis*, which uses the CBB cycle for DIC fixation (Füssel et al. 2017) had ∼2-times lower DIC fixation yields compared to *Nitrospina* and *Nitrospira* which use a O_2_-tolerant version of the reverse TCA cycle (Lücker et al. 2010, 2013). Zhang et al. (2020) measured ∼1.7-times lower DIC fixation yields of *Nitrospina gracilis* 3/211 and a terrestrial *Nitrospira* isolate compared to values in our study. We observed that radiotracer incubations conducted over the entire length of the growth curve (until early stationary phase, see Fig. S2) resulted in ∼1.4 to 1.7-times higher DIC fixation yields of NOB compared to incubations conducted until late exponential growth (when NO_2_^-^ was completely oxidized) (Fig. 2), suggesting that, in contrast to AOA where ammonia oxidation and DIC fixation were tightly coupled, nitrite oxidation might be partly decoupled from DIC fixation in NOB. While short incubation times (24 h) are typically favored over longer times for environmental measurements to avoid cross-feeding of reaction products, our results indicate that DIC fixation yields of NOB might be underestimated using these established protocols (Fig. 2).

**Fig. 2.**
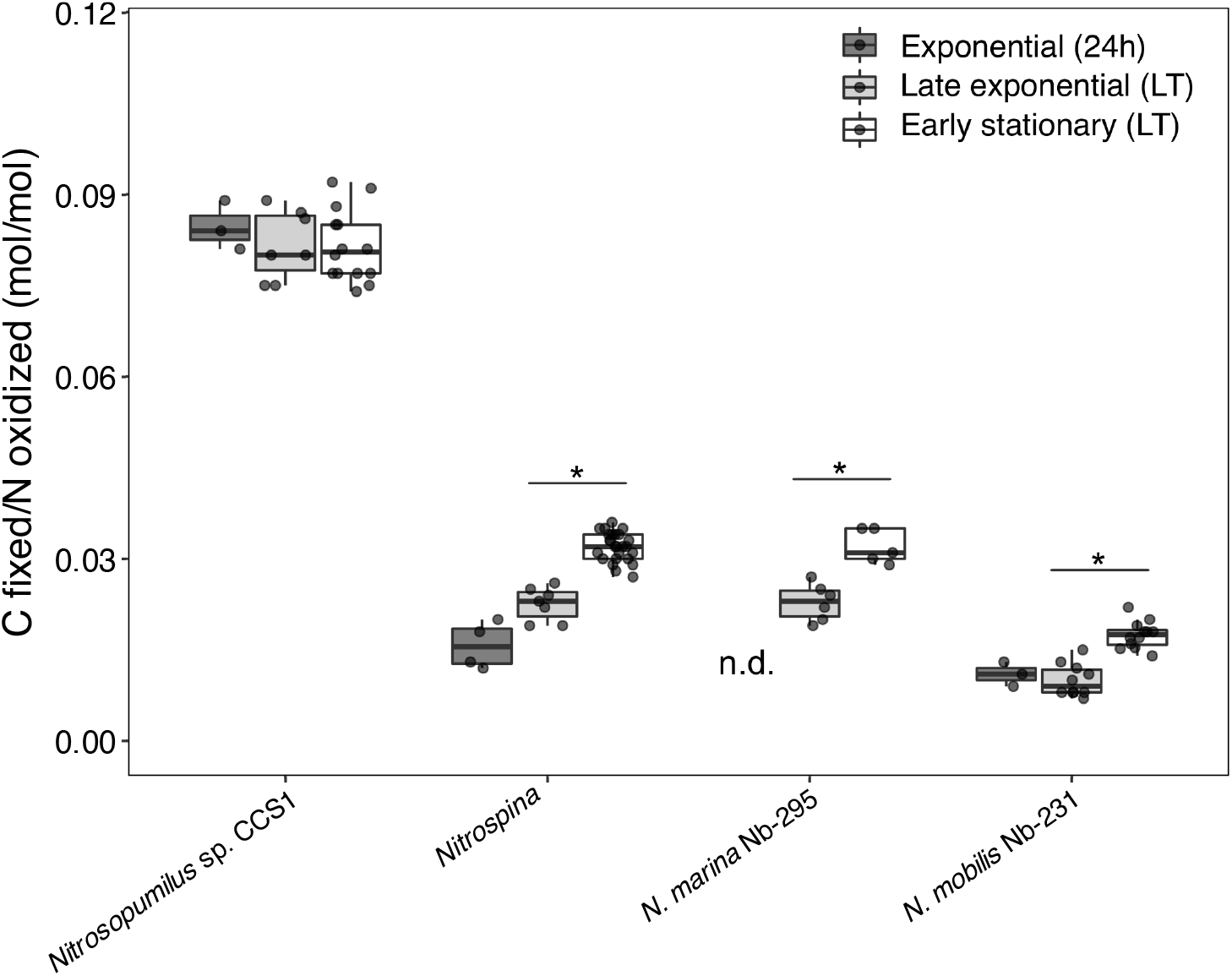
Comparison of DIC fixation yields obtained from short-term (24h) radiotracer incubations during exponential growth, and long-term (LT) radiotracer incubations carried out until either late exponential growth or early stationary phase. Measurements of both *Nitrospina* strains (Nb-3 and Nb-211) were combined in this plot. Statistical significance (adj. *p*-value <0.01) of within-condition comparisons are indicated by an asterisk (*). Statistical results of all pairwise comparisons are reported in Table S2. Representative growth curves can be found in the Supporting Information (Fig. S2).

We further explored the effect of multiple culture conditions, including environmentally relevant conditions of low substrate concentrations (1 µM) and low temperature (15°C), on DIC fixation yields of *Nitrosopumilus* sp. CCS1 and *Nitrospina* sp. Nb-3. We observed that *Nitrospina* sp. Nb-3 was ∼1.4-times more efficient in converting energy to growth when grown in natural seawater compared to artificial seawater medium, which was not observed for *Nitrosopumilus* sp. CCS1 (Fig. 3). We hypothesize that reduced N compounds present in natural seawater (ammonium and/or organic N compounds) might be responsible for the observed differences due to the high metabolic costs (6 reducing equivalents) associated with assimilatory NO_2_^-^ reduction (Einsle et al. 2002). Additions of ammonium or tryptone to artificial seawater medium likewise resulted in significantly higher DIC fixation yields (Fig. 3, Fig. S3), corroborating this hypothesis. Environmental populations of *Nitrospinae* have previously been shown to favor ammonium and the organic N sources urea and cyanate over nitrite (Kitzinger et al. 2020). Our data suggest that in addition to urea and cyanate, marine NOB can assimilate more complex organic N sources such as peptides and/or amino acids thereby saving energy that can instead be invested in C assimilation. The two most recent estimates for global ocean DIC fixation by NOB differ by one order of magnitude (Pachiadaki et al. 2017; Zhang et al. 2020) (Table S3), potentially also reflecting some of these uncertainties. Furthermore, we observed slightly higher DIC fixation yields of *Nitrosopumilus* sp. CCS1 in HEPES-buffered artificial seawater compared to unbuffered culture medium (Fig. 3), which coincided with higher cellular C quota (Table 1). While we cannot explain these observations, the differences in DIC fixation yield did not seem to be caused by variations in pH, which remained constant in unbuffered culture medium containing low substrate concentrations (1 µM NH_4_^+^).

**Fig. 3.**
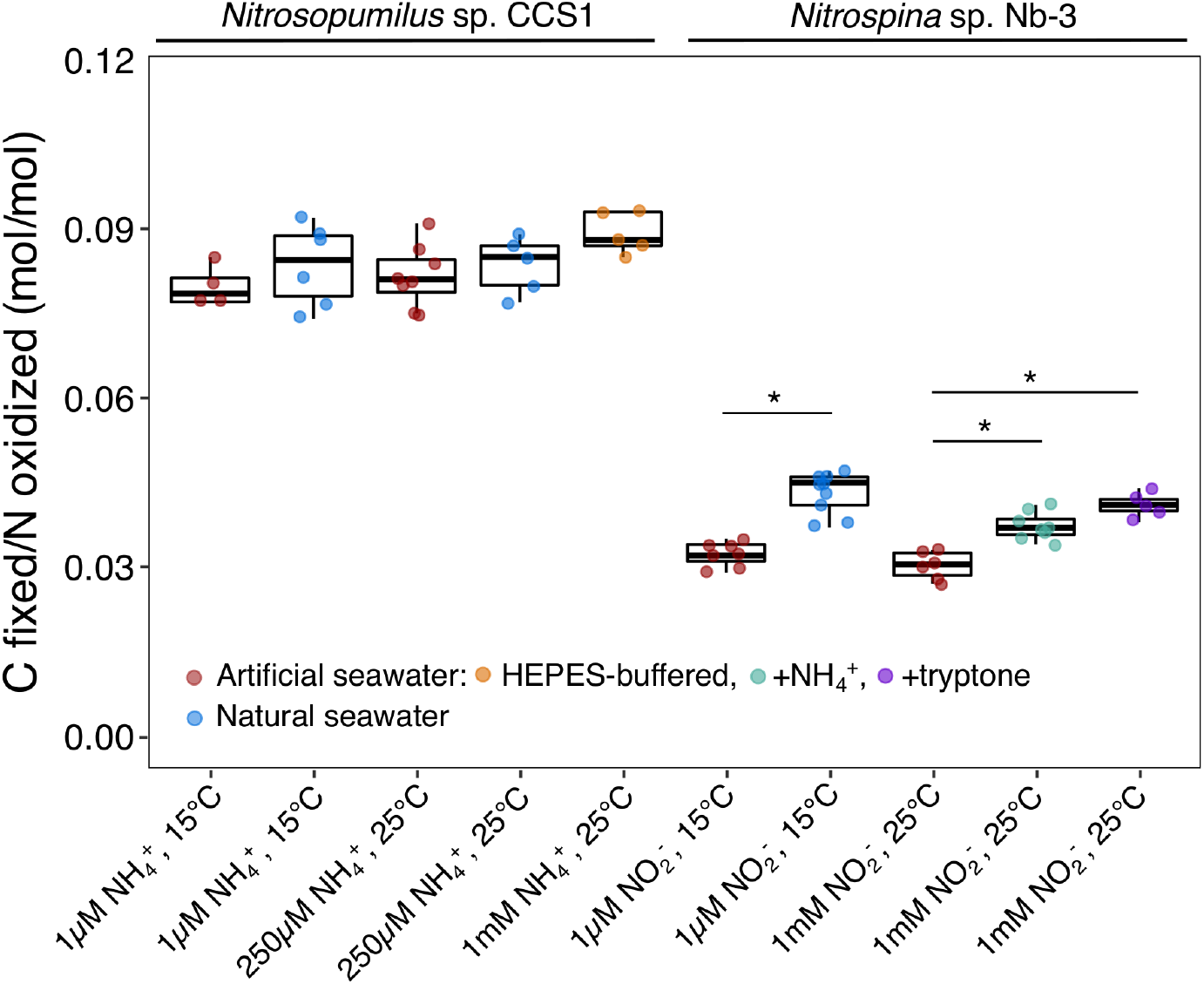
DIC fixation yields of *Nitrosopumilus* sp. CCS1 and *Nitrospina* sp. Nb-3 under different culture conditions (substrate concentrations: 1µM, 250µM, 1mM; temperature: 15°C, 25°C) and culture media (natural seawater, artificial seawater, HEPES-buffered artificial seawater). Plotted values include both, the fraction of C incorporated into biomass and the fraction of C released as DOC. Ammonium (50µM) or tryptone (150 mg L^−1^) served as additional, reduced nitrogen source for *Nitrospina* sp. Nb-3. Statistical significance (adj. *p*-value <0.01) of within-condition comparisons are indicated by an asterisk (*). Statistical results of pairwise comparisons are reported in Table S2.

The theoretical Gibbs free energy release (ΔG) for conditions in our study was 3.6-times higher for ammonia compared to nitrite oxidation (Table 2), yet DIC fixation yields of *Nitrosopumilus* sp. CCS1 and *Ca*. Nitrosopelagicus brevis U25 (Table 2) were only 2 to 2.6-times higher compared to *Nitrospina* sp. Nb-3. Similar observations were made by Kitzinger et al (2020) who reported that Nitrospinae bacteria in low O_2_ waters of the Gulf of Mexico are more efficient in translating energy gained from nitrite to C assimilation than AOA are in translating energy gained from ammonia oxidation. In addition to thermodynamics and the efficiency of the DIC fixation pathway itself, additional factors can contribute to realized energy yields, including the requirement of four out of six generated electrons by ammonia monooxygenase to reduce molecular oxygen in ammonia oxidizers (Stahl and de la Torre 2012; Caranto and Lancaster 2018). When considering that a maximum of 53.8% of the energy released from catabolism are available to ammonia oxidizers for growth (González-Cabaleiro et al. 2019), AOA appear to have slightly higher DIC fixation efficiencies compared to NOB encoding the rTCA cycle (Table 2). While oxygen protection likely increases the energy demands of the rTCA cycle (Berg 2011), our results indicate that the cycle is also highly efficient under oxic conditions that are found in most regions of the global ocean.

**Table 2.** Thermodynamic considerations and comparison of DIC fixation efficiencies and biomass yields of marine AOA and NOB grown under environmentally relevant conditions (substrate concentration: 1µM; temperature: 15°C) in artificial and natural seawater medium. Gibbs free energy calculations for NH_3_ oxidation and NO_2_^-^ oxidation can be found in Table S1.

|  | *Ca. Nitrosopelagicus* U25<sup>§</sup> | *Nitrosopumilus* sp. CCS1 |  | *Nitrospina* sp. Nb-3 |  |
| --- | --- | --- | --- | --- | --- |
| Culture medium | Natural seawater | Artificial seawater | Natural seawater | Artificial seawater | Natural seawater |
| **Gibbs free energy (kJ mol<sup>-1</sup>)** | 280 / 151* | 276 / 149* | 276 / 149* | 77 | 77 |
| **DIC fixation yield (mol mol<sup>-1</sup>)** | 0.111 ± 0.008 | 0.080 ± 0.004 | 0.085 ± 0.008 | 0.032 ± 0.002 | 0.043 ± 0.004 |
| **DIC fixation efficiency (μmol C kJ<sup>-1</sup>)** | 396 ± 29 / 735 ± 53* | 290 ± 15 / 537 ± 27* | 308 ± 29 / 570 ± 54* | 416 ± 26 | 558 ± 52 |
| **Biomass yield<sup>&</sup> (gBio gN<sup>-1</sup>)** | 0.187 ± 0.019 | 0.135 ± 0.010 | 0.143 ± 0.019 | 0.056 ± 0.005 | 0.076 ± 0.010 |

<sup>§</sup>*Ca. Nitrosopelagicus* U25 was grown at 22°C with initial substrate concentrations of 50 μM.
<sup>&</sup>The average chemical formula of bacterial biomass (CH<sub>1.7</sub>O<sub>0.4</sub>N<sub>0.2</sub>, (Popovic 2019)) was adjusted using the C:N ratios from Table 1 (AOA: CH<sub>1.7</sub>O<sub>0.4</sub>N<sub>0.25</sub>; *Nitrospina*: CH<sub>1.7</sub>O<sub>0.4</sub>N<sub>0.29</sub>).
\*When considering 53.8% of the energy released is available for growth according to González-Cabaleiro et al. 2019.

Multiple studies have used estimates of DIC fixation yields to infer DIC fixation rates associated with nitrification in diverse marine and estuarine environments (Dore and Karl 1996; Lam et al. 2004; Wuchter et al. 2006; Middelburg 2011; Lee et al. 2015), and a value of 0.1 for archaeal ammonia oxidation has widely been used in the literature (Wuchter et al. 2006; Reinthaler et al. 2010; Middelburg 2011) without direct experimental evidence. Previous measurements of DIC fixation yields were mainly derived from cultures of ammonia and nitrite oxidizers that are not representative for the majority of nitrifiers found in marine environments and were highly variable (AOB: 0.033-0.130; NOB: 0.013-0.031; (Prosser 1990) *and references therein*). The variations in DIC fixation yields we observe for marine nitrifiers across different species and culture conditions are comparably low within AOA (mean±sd=0.091 ± 0.012; *n*=47) and *Nitrospina/Nitrospira* (mean±sd=0.036 ± 0.005; *n*=56), suggesting that these values are more constrained than previous estimates and particularly useful for modelling approaches in marine systems.

### DOC release by chemolithoautotophs

We measured DOC release rates of ten nitrifier cultures and tested how different culture conditions affected the amount of DOC released in proportion to the amount of fixed DIC. All investigated strains released DOC during exponential growth, and DOC release ceased when cultures reached stationary phase (as determined by comparing the total amount of released DOC until late exponential vs stationary phase, see Fig. S4), suggesting that DOC release is a feature of metabolically active nitrifiers. This is in agreement with earlier observations of amino acid release by exponentially growing *Nitrosopumilus* cells (Bayer et al. 2019a). The amount of chemoautotrophically fixed DIC that was released as DOC by nitrifiers made up on average ∼5-15% (Fig. 4a). This is within the range observed for phytoplankton, which released 2-10% and 4-42% of their photosynthetically fixed DIC in culture and environmental studies, respectively (Carlson 2002), *and references therein*).

**Fig. 4.**
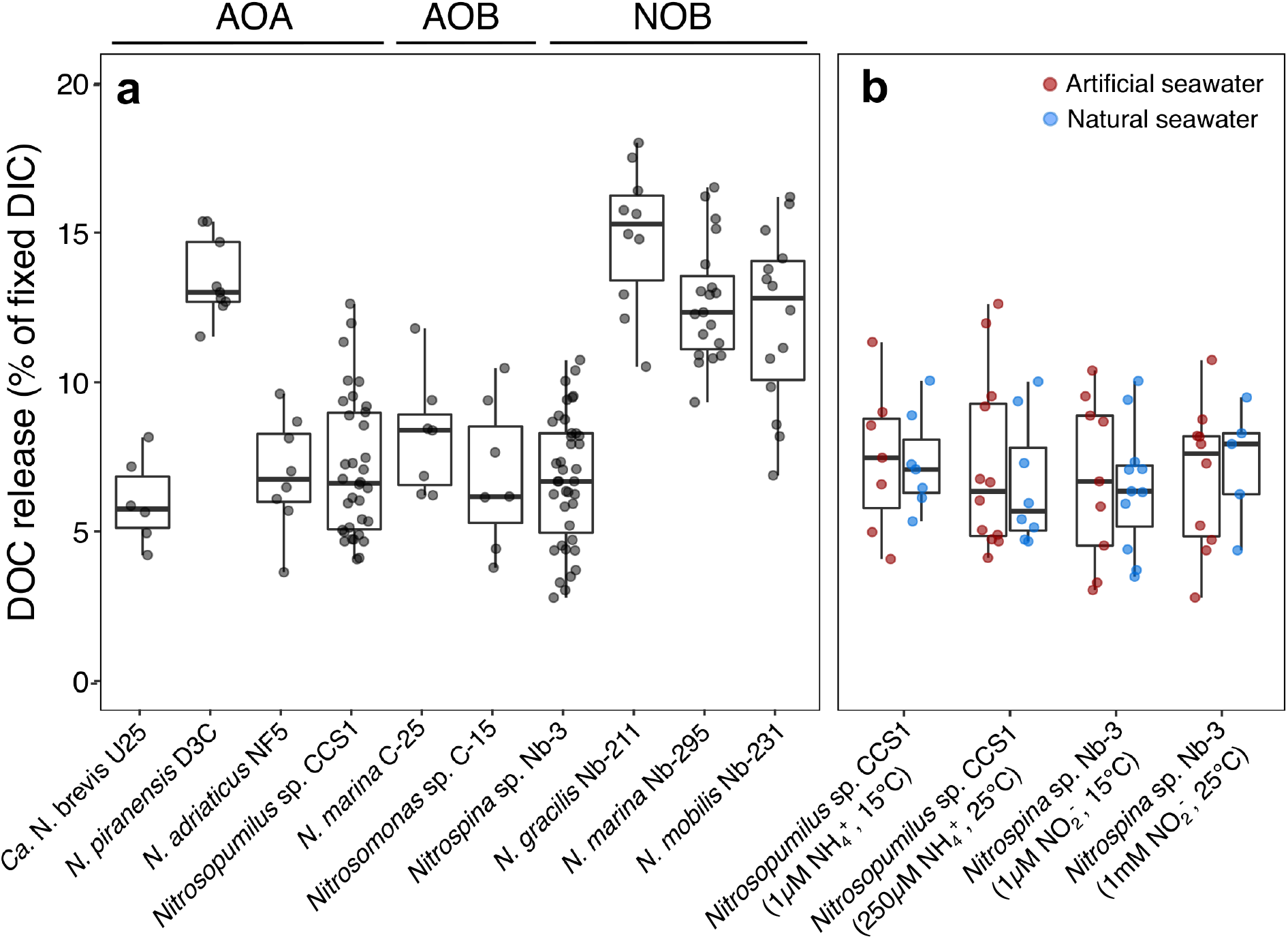
DOC release by marine nitrifiers as a fraction of fixed DIC. **a)** Comparison of DOC release by ten different phylogenetically diverse marine nitrifiers. Values obtained from cultures grown under different conditions (see panel b) are included in this plot. DOC release by *Ca*. N. brevis might be underestimated due to the presence of heterotrophic bacteria that could take up some of the released DOC. **b)** Comparison of DOC release by *Nitrosopumilus* sp. CCS1 and *Nitrospina* sp. Nb-3 grown under different culture conditions (substrate concentrations: 1µM, 250µM, 1mM; temperature: 15°C, 25°C) in artificial or natural seawater medium. Statistical results of all pairwise comparisons are reported in Table S2.

DOC release varied between closely related species (Fig. 4a). *Nitrosopumilus piranensis* released more DOC compared to the two other investigated *Nitrosopumilus* species, which is in agreement with (Bayer et al. 2019b) who reported higher amino acid release rates of *N. piranensis* compared to *N. adriaticus*. Differences in the amount of released DOC have also been recently reported between the closely related aquarium strain *Nitrosopumilus maritimus* SCM1 (9-19% of fixed DIC) and the environmental strain *Nitrosopumilus maritimus* NAOA6 (5% of fixed DIC) (Meador et al. 2020). Within NOB, *Nitrospina* sp. Nb-3 consistently released less DOC compared to *N. gracilis* Nb-211 and the two phylogenetically more distantly related species *N. marina* and *N. mobilis*.

The fraction of released DOC remained constant across different culture conditions including environmentally relevant conditions of low substrate concentration (1 µM) and at low temperature (15°C) in natural seawater (Fig. 4b). This suggests that DOC release is not an artifact of unrealistic culture conditions but likely a feature exhibited by nitrifier populations in the environment. While the composition of DOM released by bacterial nitrifiers is currently unknown, a fraction of the DOM released by AOA has been shown to consist of labile compounds, such as amino acids, thymidine and B vitamins, that can limit microbial heterotrophic activity in open ocean waters (Bayer et al. 2019a).

## Conclusions

Our results suggest that DIC fixation yields of marine NOB might be underestimated by conventional short-term tracer incubations, due to a partial decoupling between NO_2_^-^ oxidation and C assimilation. Additionally, DIC fixation yields of *Nitrospina* were positively affected by the presence of ammonium or complex organic N compounds, which might influence metabolic interactions with ammonia oxidizers and/or heterotrophic prokaryotes in the environment, suggesting a potentially underappreciated role for competition in the N cycle (Santoro 2016).

DIC fixation yields of marine nitrifiers obtained in our study will help to further constrain the relationship between C and N fluxes in the nitrification process and inform theoretical models about how to connect observations at microscale to regional and global scales. Using a mean global value of organic C export from the euphotic zone of ∼6 Pg C yr^−1^ (Siegel et al. 2014) and a mean C:N ratio of sinking marine particles (at the surface) of ∼7.1 (Schneider et al. 2003), we estimate that the resulting global ocean organic N export of ∼0.85 Pg yr^−1^ could fuel up to 0.13 Pg C y^-1^ of chemoautotrophic DIC fixation (0.094 Pg C y^-1^ by AOA and 0.037 Pg C y^-1^ by NOB) in the dark ocean, which is lower than previous estimates (0.15-1.4 Pg C y^-1^, see Table S3 and references therein). Furthermore, we show that nitrifiers release significant amounts of DOC under environmentally relevant conditions, equating to fluxes of 0.006-0.02 Pg C y^-1^ of fixed DIC released as DOC. Elucidating the lability and fate of the DOM released by nitrifiers will be crucial to understand its implications for the marine carbon cycle.

## Acknowledgements

We thank Thomas Reinthaler for advice on DIC fixation measurements and for sharing his lab protocols, and Elisa Halewood and Janice Jones for support and advice with setting up the DIC fixation method in the lab. We also thank Sylvia Kim for total alkalinity and pH measurements, Kenneth Marchus for elemental (CHN) analyses, Christopher Sedlacek for sending culture stocks of *N. adriaticus* and *N. piranensis*, Claus Pelikan for advice on statistical analyses, and Holger Daims for advice on bioenergetic considerations. Frederica Valois and John B. Waterbury are acknowledged for their decades long commitment and maintenance of the nitrifier culture collection at the Woods Hole Oceanographic Institution. This research was supported by a Simons Foundation Early Career Investigator in Marine Microbiology and Evolution Award (345889) and a US National Science Foundation (NSF) award OCE-1924512 to AES. BB was supported by the Austrian Science Fund (FWF) Project Number: J4426-B (“The influence of nitrifiers on the oceanic carbon cycle”). Support for CAC was provided by Simons Foundation International’s BIOS-SCOPE program.

The authors declare no conflict of interest.

## Notes

### Competing Interest Statement

The authors have declared no competing interest.

