## Supporting Information for "Carbon content, carbon fixation yield and dissolved organic carbon release from diverse marine nitrifiers"

### Supporting Tables and Figures

**Table S1.** Gibbs Free Energy calculations of ammonia and nitrite oxidation (provided as separate excel sheet)

**Table S2.** Results of statistical analyses (see Materials and Methods section in the main text). **a)** Pairwise comparisons of DIC fixation yields of marine nitrifiers obtained under different growth phases (EXP, exponential; STAT, stationary) and incubation times (24h; LT, long-term). **b)** Pairwise comparison of DIC fixation yields of *Nitrosopumilus* sp. CCS1 grown in different culture medium (ASW, artificial seawater; NSW, natural seawater; ASW-HEPES, HEPES-buffered artificial seawater). **c)** Pairwise comparisons of DIC fixation yields of marine NOB grown in different culture medium (ASW, artificial seawater; NSW, natural seawater) with additions of ammonium (+$NH_4^+$) or tryptone (+trp) and at different temperature (15°C, 25°C) and $NO_2^-$ concentrations (1µM, 1mM $NO_2^-$).

**a)**

| | *Nitrosopumilus* sp. CCS1 | | *Nitrospina* sp. Nb-3 | | *N. mobilis* Nb-231 | | *N. marina* Nb-295 | |
|---|---|---|---|---|---|---|---|---|
| | EXP (24h) | EXP (LT) | EXP (24h) | EXP (LT) | EXP (24h) | EXP (LT) | EXP (24h) | EXP (LT) |
| EXP (LT) | 0.536291 | | 0.023029 | | 0.390098 | | n.d. | |
| STAT (LT) | 0.536291 | 0.94014 | **0.002234** | **0.000188** | 0.013498 | **0.000362** | n.d. | **0.006054** |

**b)**

| | *Nitrosopumilus* sp. CCS1 | |
|---|---|---|
| | ASW | ASW-HEPES |
| ASW-HEPES | 0.020935 | |
| NSW | 0.27895 | 0.166714 |

**c)**

| | *Nitrospina* sp. Nb-3 | | | | *N. marina* Nb-295 |
|---|---|---|---|---|---|
| | ASW (1µM, 15°C) | NSW (1µM, 15°C) | ASW (1mM, 25°C) | ASW+$NH_4^+$ (1mM, 25°C) | ASW (1mM, 25°C) |
| NSW (1µM, 15°C) | **0.006338** | | | | n.d. |
| ASW (1mM, 25°C) | 0.19151 | **0.006338** | | | |
| ASW+$NH_4^+$ (1mM, 25°C) | **0.007453** | **0.008404** | **0.006338** | | n.d. |
| ASW+trp (1mM, 25°C) | **0.008404** | 0.203277 | **0.008648** | 0.028146 | **0.003184** |

**Table S3.** Comparison of estimates of global dark ocean DIC fixation fueled by ammonia and nitrite oxidation

| **Nitrification-fueled global dark ocean DIC fixation ($Pg\ C\ y^{-1}$)** | **Reference** |
|---|---|
| 0.13 ($NH_3$ oxidation: 0.095; $NO_2^-$ oxidation: 0.037) | this study |
| 0.15 ($NH_3$ oxidation: 0.12; $NO_2^-$ oxidation: 0.031) | (Zakem et al. 2018) |
| 0.15-0.2 ($NH_3$ oxidation: 0.11-0.14; $NO_2^-$ oxidation: 0.036-0.06) | (Zhang et al. 2020) |
| 0.11 ($NH_3$ oxidation only) | (Middelburg 2011) |
| 0.4 ($NH_3$ oxidation only) | (Herndl et al. 2005; Wuchter et al. 2006) |
| 1 ($NO_2^-$ oxidation only) | (Pachiadaki et al. 2017) |

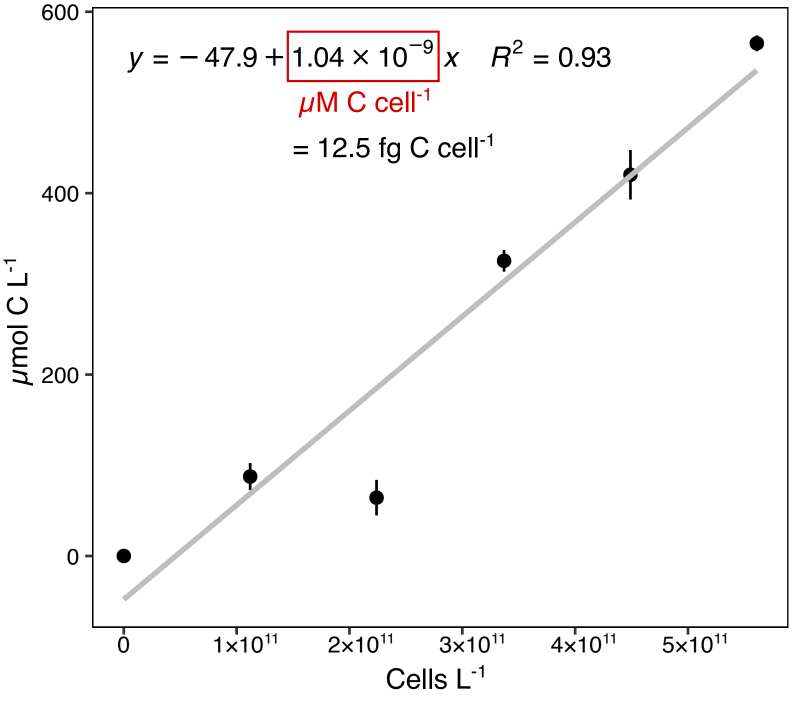


**Fig. S1** Dilution series of concentrated cells to determine the cellular C content of *Nitrosopumilus* sp. CCS1

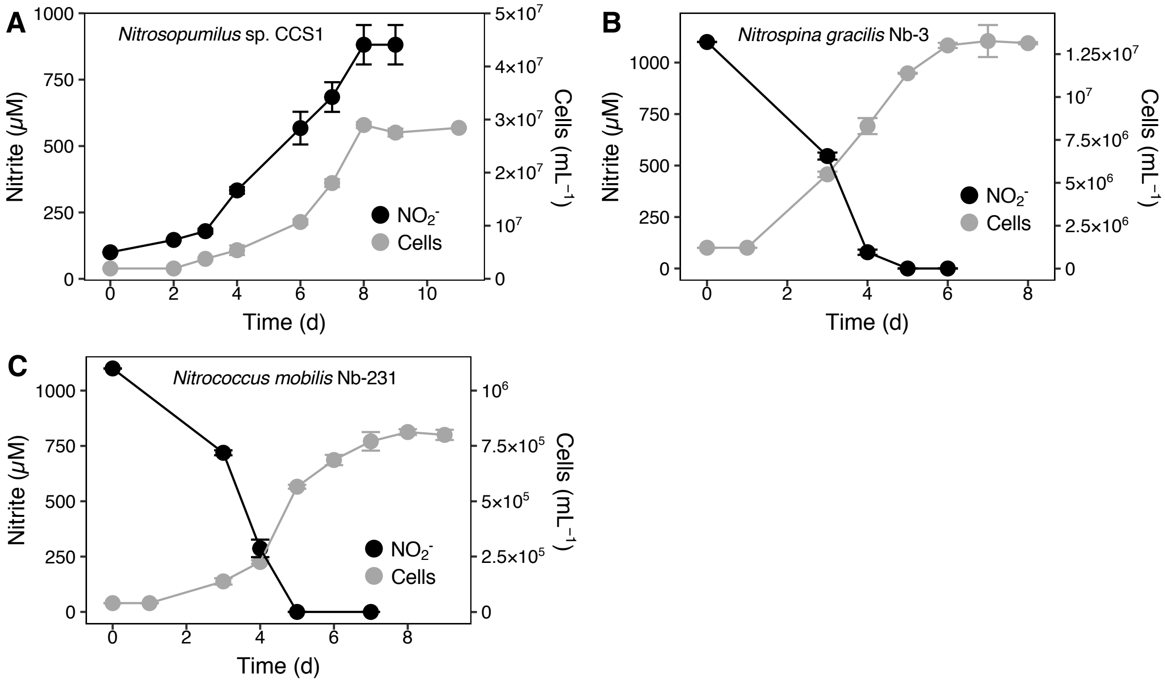


**Fig. S2** Nitrite and cell concentrations of *Nitrosopumilus* sp. CCS1 (A), *Nitrospina* sp. Nb-3 (B) and *Nitrococcus mobilis* Nb-231 (C).

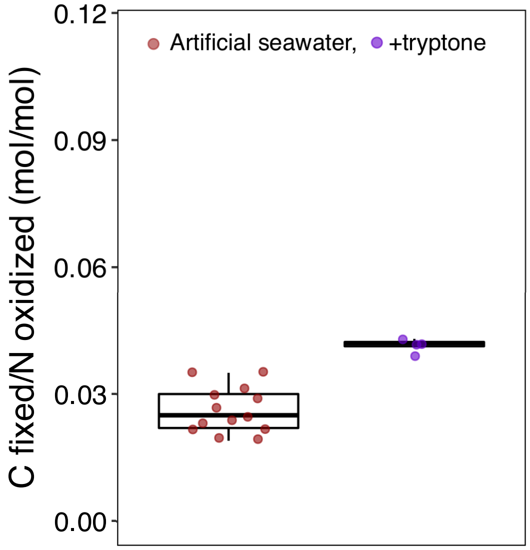


**Fig. S3.** DIC fixation yield of *N. marina* Nb-295 grown in artificial seawater medium with and without addition of tryptone.

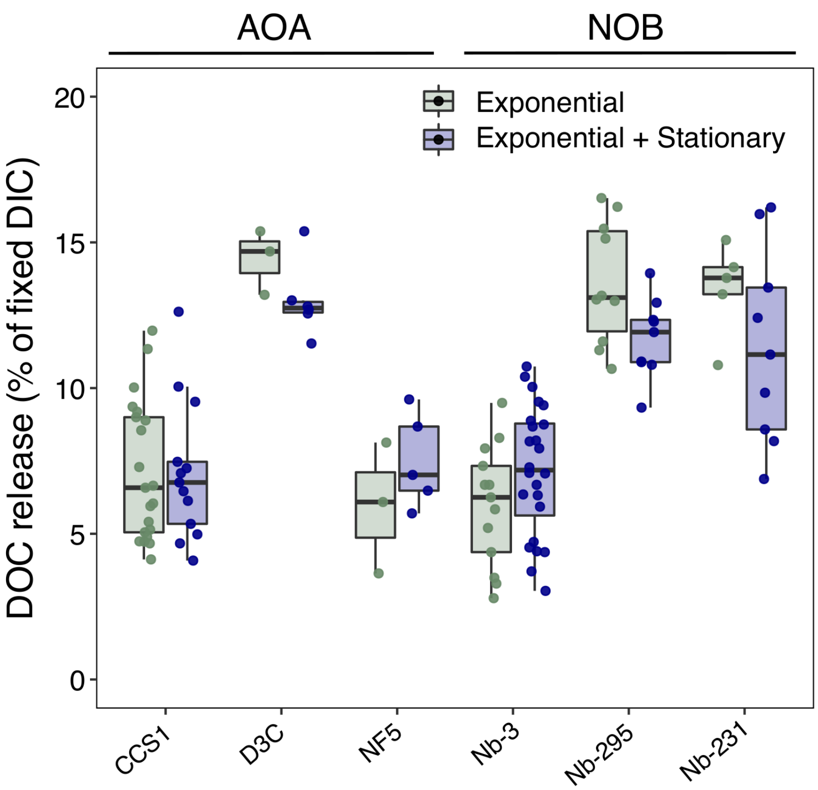


**Fig. S4.** DOC release (as a proportion of fixed DIC) of marine nitrifiers during different growth phases.
